# Conserved features of eye movement related eardrum oscillations (EMREOs) across humans and monkeys

**DOI:** 10.1101/2023.03.08.531768

**Authors:** Stephanie N Lovich, Cynthia D King, David L.K. Murphy, Hossein Abbasi, Patrick Bruns, Christopher A Shera, Jennifer Groh

**Affiliations:** Department of Psychology and Neuroscience, Duke University; Department of Neurobiology, Duke University; Center for Cognitive Neuroscience, Duke University; Duke Institute for Brain Sciences, Duke University; Department of Psychiatry and Behavioral Sciences, Duke University; Biological Psychology and Neuropsychology, University of Hamburg; Department of Otolaryngology, University of Southern California; Department of Computer Science, Duke University; Department of Biomedical Engineering, Duke University

**Keywords:** Visual-auditory integration, eye movements, reference frame, otoacoustic emissions, middle ear muscles, EMREO

## Abstract

Auditory and visual information involve different coordinate systems, with auditory spatial cues anchored to the head and visual spatial cues anchored to the eyes. Information about eye movements is therefore critical for reconciling visual and auditory spatial signals. The recent discovery of eye movement-related eardrum oscillations (EMREOs) suggests that this process could begin as early as the auditory periphery. How this reconciliation might happen remains poorly understood. Because humans and monkeys both have mobile eyes and therefore both must perform this shift of reference frames, comparison of the EMREO across species can provide insights to shared and therefore important parameters of the signal. Here we show that rhesus monkeys, like humans, have a consistent, significant EMREO signal that carries parametric information about eye displacement as well as onset times of eye movements. The dependence of the EMREO on the horizontal displacement of the eye is its most consistent feature, and is shared across behavioral tasks, subjects, and species. Differences chiefly involve the waveform frequency (higher in monkeys than in humans) and patterns of individual variation (more prominent in monkeys than humans), and the waveform of the EMREO when factors due to horizontal and vertical eye displacements were controlled for.

## Introduction

Linking visual and auditory space requires information about eye movements. Spatial cues to sound location involve interaural level differences (ILDs) and interaural timing differences (ITDs), cues that can be used to determine the positions of sounds with respect to the head, or a head-centered reference frame. In contrast, the locations of visual stimuli are detected via the pattern of light on the retina, an eye-centered reference frame. In short, every time the eyes move, the retina shifts with respect to the head and ears, changing the relationship between eye-centered visual spatial cues and head-centered auditory spatial cues. Thus, the brain must keep track of eye movements to resolve the difference between these two coordinate systems and support binding of visual and auditory stimuli based on their location (e.g.(Groh and Sparks, 1992; Groh, 2014)).

Early work on how the brain achieves reference frame alignment was conducted via single unit recordings in rhesus monkeys. In the superior colliculus, changes in eye position were found to cause auditory receptive fields to shift their positions with respect to the head (Jay and Sparks, 1984, 1987a; Lee and Groh, 2012); (see also Hartline et al., 1995; Zella et al., 2001; Populin et al., 2004). This suggested that a coordinate transformation of auditory signals was occurring somewhere within the brain (Groh and Sparks, 1992). Subsequent work extended these observations to other multisensory brain regions such as the frontal eye fields (Russo and Bruce, 1994; Caruso et al., 2019) and parietal cortex (Stricanne et al., 1996; Cohen and Andersen, 2000; Mullette-Gillman et al., 2005, 2009). Intriguingly, effects of eye position were also found in predominantly auditory brain regions such as auditory cortex (Werner-Reiss et al., 2003; Fu et al., 2004; Maier and Groh, 2010) and the inferior colliculus (Groh et al., 2001; Zwiers et al., 2004; Porter et al., 2006; Bulkin and Groh, 2012a, b; Willett et al., 2019) during responses to sound stimuli.

These reports concerning eye movement effects on auditory responses in auditory brain regions motivated our foray into the most peripheral part of the auditory system, the ear itself. We reasoned that information about eye movements could be conveyed via the descending pathways to the motor actuators within the ear, such as the middle ear muscles and the outer hair cells, and that, just like with conventional otoacoustic emissions and middle ear reflex testing (e.g. Kemp, 1978; Shera, 2004; Schairer et al., 2013), the impact of such signals might produce movements of the eardrum that could be detected by microphones in the ear canal. This led to the discovery of eye movement-related eardrum oscillations (EMREOs) (Gruters et al., 2018; Murphy et al., 2020; Lovich et al., 2022; King et al., 2023) and inquiries into their relationship to visual and auditory perception (Abbasi et al., 2023; Bröhl and Kayser, 2023).

EMREOs occur with saccades, time-locked to the beginning of, or sometimes slightly preceding, their onset, and showing phase-resetting at saccade offset. Then oscillation then continues for at least several tens of milliseconds after the eyes stop moving. The waveforms of EMREOs carry precise and parametric information about the direction and amplitude of the saccade (Lovich et al., 2022). EMREOs occur in the absence of incoming sound, and how they (or more properly, the underlying mechanism they reflect) impact sound transduction is unknown. It is also unknown how they might contribute to the eye movement-related modulation of sound responses observed later in the brain’s auditory pathway (Russo and Bruce, 1994; Caruso et al., 2019) (Stricanne et al., 1996; Cohen and Andersen, 2000; Groh et al., 2001; Werner-Reiss et al., 2003; Fu et al., 2004; Zwiers et al., 2004; Mullette-Gillman et al., 2005; Porter et al., 2006; Mullette-Gillman et al., 2009; Maier and Groh, 2010; Bulkin and Groh, 2012a, b; Willett et al., 2019).

Intriguingly, EMREOs are seen in both humans and rhesus monkeys (Gruters et al., 2018). A detailed comparison between the human and monkey EMREOs can therefore serve as a natural experiment to shed light on which features of the EMREO are most likely to be functionally important to auditory coordinate transformations or any other potential purpose. This human-monkey comparison is particularly apt because the two species have similar visual and auditory acuity and similar eye movements (Jay and Sparks, 1990). They are also known to integrate visual and auditory space in reasonably similar ways, showing similar thresholds for fusing vs. distinguishing visual and auditory locations (Mohl et al., 2019). Features of the EMREO that are shared across humans and monkeys may represent aspects that are particularly important for these shared perceptual and oculomotor attributes. In short, a quantitative cross-species comparative analysis can shed light on the function and underlying mechanism of this phenomenon, and is a needed advance over the qualitative assessment provided in our initial study (Gruters et al., 2018).

We report here that EMREOs in both humans and monkeys exhibit a similar dependence on the horizontal displacement of the eye during the accompanying saccade. Differences across species include the frequency of the EMREO waveform, which is higher in monkeys than in humans, and the dependence of the EMREO on the vertical displacement of the eye, which showed more variation across individual monkeys than in individual humans, causing this signal to “wash out” when averaged across monkeys. Monkeys and humans also differed in the nature of the EMREO waveform when the dependence on horizontal and vertical saccade amplitude is controlled for.

However, EMREOs were similar across different task designs in humans, suggesting that they are associated with the eye movements regardless of how the eye movements are elicited. We hypothesize that the eye displacement signals reflected in EMREOs may be used during auditory localization and perception, and that the horizontal dependence may be specifically used to tailor interaural timing and level difference cues to sound location in an eye movement dependent fashion.

## Methods

### Participants

We recorded eye movement-related eardrum oscillations (EMREOs) simultaneously in the left and right ears of 4 monkeys (3 female, 1 male) and 21 human subjects (14 female, 7 male). All procedures involving human subjects were approved by the Duke University Institutional Review Board. All procedures involving monkey subjects were performed in accordance with an animal protocol approved by Duke University IACUC. Subjects had apparently normal hearing and normal or corrected vision. Informed consent was obtained from all human participants before testing, and all human participants received monetary compensation for participation.

### Tasks

#### Monkey behavioral task

To maximize the rate of data collection in the monkeys and permit the testing of relatively untrained animals, we allowed monkeys to freely view a black monitor screen and make saccades wherever and whenever they wished (free-viewing saccade task). The black screen made capturing the pupil of the monkey easier and more accurate because of size of the pupil in darkness. The free viewing task allowed us to test any of the available monkeys in our colony regardless of their training status. No external sounds were presented.

#### Human behavioral tasks

Since our previous study involved performance of visually guided saccades to specific targets (Gruters, Murphy et al. 2018), but the monkey testing for the present study involved free viewing, we tested 4 of our human subjects on both a visually-guided saccade task similar to the one we used before (Figure 1a) and a free-viewing task (Figure 1b) similar to the one used for monkeys (Figure 1c). The remainder of the human subjects were tested only on the visually-guided saccade task. For the free-viewing task, humans viewed abstract Jackson Pollock paintings, which helped them stay awake and engaged throughout the task. They made saccades wherever and whenever they wished, similar to the monkeys. This permitted us to establish whether different tasks affect the EMREO signals (in humans) and to compare the monkey and human results using a free-viewing task. No external sounds were presented during either task.

**Figure 1:**
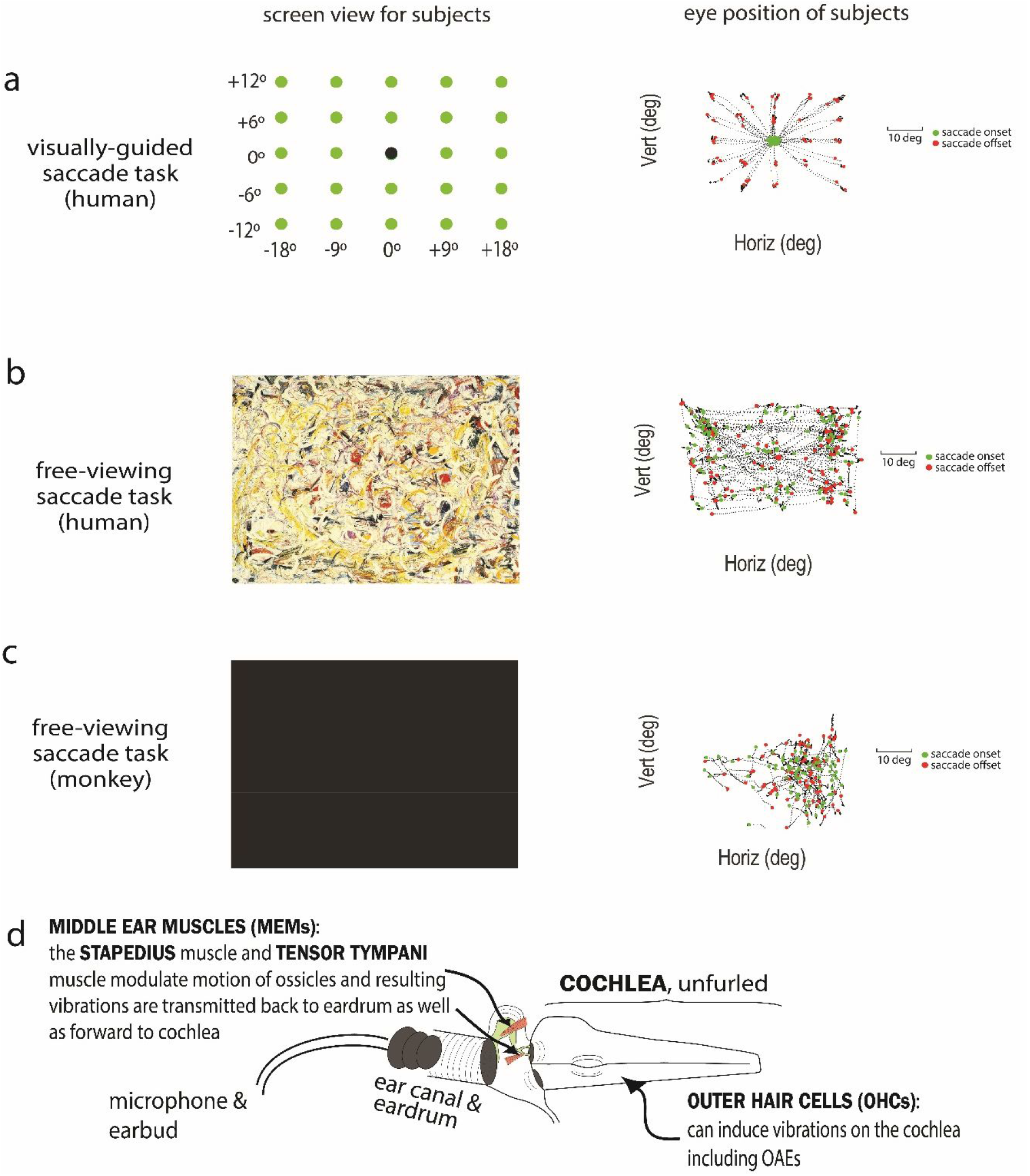
Task and microphone setup. 1a. Humans performed a visually-guided saccade task in which the black fixation dot appeared on the screen for 750 ms. The black dot would then disappear and one of the green dots would appear for 750 ms. Subjects had to saccade to the green dot and fixate for a minimum of 200 ms at which point the dot would turn red signaling that the trial was over. An example of one subject’s eye position during one block can be seen on the right. 1b. Humans performed a free-viewing saccade task interleaved with the visually-guided saccade task in which they made saccades in any direction and magnitude of their own volition. An example of one subject’s eye position during one block can be seen on the right. 1c. Monkeys performed the same free-viewing task as humans but with a black screen. An example of one monkey’s eye position during one block can be seen on the right. 1d. A microphone was placed into each ear canal of the subject and recorded pressure changes of the eardrum while the subject made saccades. No external sounds were presented.

### Sessions, Head Restraint, Eye Tracking, and Saccade identification

#### Sessions

An earphone assembly consisting of a microphone (Etymotic ER-10B+) and transducer (Etymotic ER 2) was placed in each ear of the subject while they made saccades (figure 1d). In a given session, human subjects performed 3 blocks (5 minutes/block) of the visually-guided saccade task interleaved with similar length blocks of free viewing, for a total of six blocks. Each session lasted approximately one hour. Microphones (48 kHz sampling rate) were assessed prior to each session by placing each earphone assembly into a test tube and playing a frequency sweep from 10Hz to 2kHz through the transducer (pure tone sweep from 10-1600Hz: 10Hz steps for 10-200Hz, 100Hz steps for 300-1000Hz, 200Hz steps for 1200-2000Hz) to evaluate recording fidelity over a broad frequency range. Microphones were also assessed before every block of a session using the same frequency sweep while in the ears of the subject. Monkey sessions lasted about 60-180 minutes. Human participants were tested in a total of 3-5 sessions each and monkeys were tested in 20-30 sessions.

#### Head restraint and eye tracking

Head movements were minimized in human participants with a chin rest and in monkey participants using a surgically implanted head post. Surgical implantation was accomplished under general anesthesia, aseptic conditions, and with suitable analgesics to minimize discomfort in accordance with IACUC regulations. Eye movements were monitored in both humans and monkeys using the same eye tracking system (EyeLink1000), and using similar brightness, pupil size, corneal reflections settings. Eye position was sampled at 1000 Hz. In most subjects, the left eye was tracked and the right eye was assumed to move in a conjugate fashion. In human subjects, the eye tracking system was calibrated based on saccades to visual targets. In the monkeys, a visual task-based calibration was not performed but the EyeLink’s base settings for gain and offset based on subject distance were used and found to be effective. We verified this by checking the monkey eye data against the following assumptions: (a) the average eye position should be roughly straight ahead, and (b) the standard deviation of the distribution of eye positions should be roughly 10 degrees. Operationally, such assumptions have worked well for us in the past when calibrating eye tracking in monkeys prior to training on visual tasks. The data generally accorded well with the standard deviation assumption, indicating the gain setting was roughly appropriate, but the center of the distributions was not necessarily centered at straight ahead for all monkeys and sessions, indicating that the offset setting was not always perfect. As described further below, the data analysis methods used were robust to differences in these calibration procedures.

### Initial analysis steps for microphone and eye tracking data

After collection, microphone data were downsampled from 48 kHz to 2 kHz for further analysis. These data were otherwise unfiltered. Saccade onsets and offsets were identified based on the rate of change in eye acceleration (third derivative, or “jerk”). After smoothing with a 7 point filter (7 ms), the first peak of the jerk was considered to be saccade onset and the second peak of the jerk was considered to be saccade offset (Murphy et al., 2020) (Lovich et al., 2022).

### Data inclusion criteria and normalization of signals across subjects/sessions

Data could be excluded at the session/block level or at the trial/saccade level. At the session/block level, data for the entire session/block were excluded if the assessment of the microphone via frequency sweep for that session or block indicated any major issues, such as a possible shift in the position of the earphone assembly. Within each session and block, individual trials or saccades could be excluded on the basis of the eye tracking data.

Trials/saccades were excluded if there was less than 200 ms of steady fixation before saccade onset or after saccade offset. Saccades could also be excluded if the saccade curvature was too great (more than 4.5º of subtended angle). These screens typically caused the exclusion of many eye movements in the free viewing tasks as they proved to be drifting movements rather than true saccades or they were not preceded and followed by satisfactory periods of steady fixation. After these screens were applied, we also excluded individual trials based on whether the RMS noise level on individual trials exceeded a criterion value (0.02, arbitrary units, set based on inspection of the normal RMS range in quiet recording sessions). We also excluded trials if the maximum or minimum microphone value on that trial was more than 10 standard deviations from the mean microphone value in that session or block. Overall, these exclusion criteria served to minimize the inclusion of trials in which the monkey generated noise via fidgeting or other behaviors. The same criteria were applied to both human and monkey sessions. For monkeys, between 7 and 37% of potential saccades were included for analysis, yielding between 9,000 and 51,000 total saccades for each subject. For the four human subjects tested similarly to the monkeys, between 24 and 86% of potential saccades were included for analysis, yielding between 700 and 1500 saccades for each subject and task condition. After these exclusion criteria were applied, we then computed the Z-scores of the microphone values relative to a pre-saccadic baseline (−100 to -40 ms prior to saccade onset). Results in the present study are therefore expressed in units of standard deviation relative to this baseline period.

### Data Analysis

Extending/replicating our previous methods (Gruters et al., 2018; Lovich et al., 2022), we fit the raw EMREO signal collected in each ear to a linear regression model that includes the basic parameters of each eye movement:

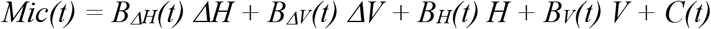

The first four terms concern the change in eye position relative to the starting position for each saccade in the horizontal and vertical dimensions, and are measured in degrees. Specifically, these terms concern the horizontal displacement of the eye (*ΔH)*, vertical displacement of the eye (*ΔV)*, the horizontal initial position (H), vertical initial position of the eye (V). Time-varying coefficients *B*_*ΔH*_*(t), B*_*ΔV*_*(t), B*_*H*_*(t)*, and *B*_*V*_*(t)* were fit to these parameters. The final term is a time-varying constant component (C(t)) that captures everything else left in the signal. C(t) can be thought of as the best fitting average oscillation across all eye positions and displacements.

This regression model allows us to analyze the EMREO with respect to the individual parameters of the synchronous eye movement and is largely robust to the particular task involved, except that the initial position parameters will mainly capture fixational scatter in the visually-guided saccade task versus a much larger range of initial positions in the free viewing tasks. In addition, as noted above, the monkey calibration procedures left the precise values of position somewhat uncertain. Thus, we do not necessarily expect perfect correspondence of the initial position coefficients of the regression fits across these two tasks; any small differences in the accuracy of eye tracking calibration will also affect the precise values of these coefficients. Accordingly, we will focus on the regression coefficients for displacement in the horizontal and vertical dimensions as well as the constant component when comparing humans and monkeys, building on our related work (Lovich et al., 2022; King et al., 2023).

## Results

In the human participants, we first confirmed that task performance had little impact on key aspects of the EMREO signal. We found that the regression coefficients of horizontal displacement (*ΔH)*, vertical displacement (*ΔV)* and the constant component (C) across four human subjects while they performed the visually-guided saccade task (purple) and the free-viewing saccade task (green) were similar (Figure 2a, data from 4 right ears). While different subjects showed different oscillatory patterns, the overall pattern was generally very consistent within subjects across tasks. The thick parts of the EMREO signal traces in Figure 2a indicate the time periods when the corresponding regression coefficient differed significantly from zero with 95% confidence, and the shaded areas are the SEM. The horizontal displacement regression coefficient (Figure 2a, top row) generally differed from zero for several tens of milliseconds starting at saccade onset except where it crossed zero as it transitioned between a peak or a trough, but there was little notable difference across tasks. The vertical term showed more variation across subjects (Figure 2a middle row) but, when present, was still quite similar within each subject for each task (e.g. S88). The constant component was also variable across subjects, but similar across tasks. Overall, these results indicate that an oscillatory dependence on horizontal saccade amplitude occurs across all subjects and for both visually-guided and free-viewing saccades, with some variation in the timing, amplitude, precise waveform, and presence/magnitude of the vertical dependence and constant components. This pattern is consistent with our previous work (Gruters et al., 2018; Murphy et al., 2020; Lovich et al., 2022; King et al., submitted), and suggests that the free-viewing saccade task is an equally effective way to collect EMREO data.

**Figure 2:**
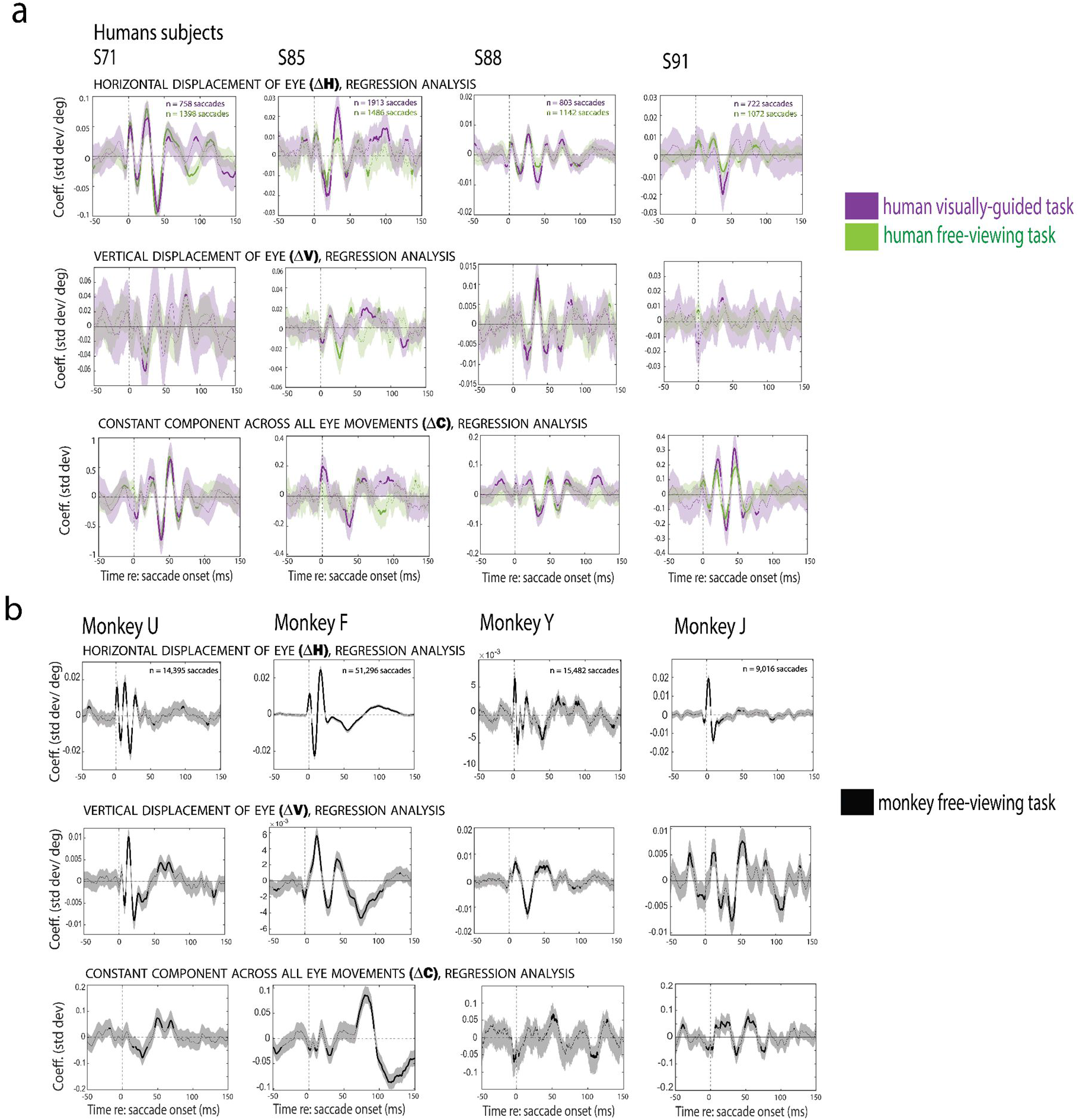
Regression analysis and individual subject data. 2a. The regression coefficient reflecting the influence of horizontal and vertical displacements of the eye on the EMREO signal (top, middle) and the constant component (bottom) in the right ears of humans across both tasks (visually-guided in purple and free viewing in green). The number of saccades for each subject/task type are noted in the horizontal component graph for each subject. 2b. Similar to (a) but for the right ears of monkeys during the free-viewing task. Shading indicates standard error of the measurement (SEM). Because there are many more saccades for the monkeys than the humans, the SEM appears smaller for this species.

Figure 2b shows corresponding findings for the right ears of four monkey subjects while they performed the free-viewing saccade task (black). Like the human data, the EMREO is strongly dependent on the horizontal displacement of the associated eye movement (Figure 2b top row). The oscillation co-varies in phase and amplitude similarly to the signal recorded in humans. Like the humans, the waveform varied to some extent across subjects, but all showed a sharp onset at (or slightly before) saccade onset, continuing for at least one full oscillatory cycle (i.e. including one peak and one trough), and all subjects had a significant signal at some points in the first few tens of milliseconds after saccade onset. Similar to the human data, the vertical and constant components varied across monkey subjects more than did the horizontal component. The vertical coefficients (middle row) differed considerably across monkeys in magnitude and waveform. As can be seen from the differences in the scales of the y-axes of the corresponding horizontal and vertical plots for each monkey, the vertical dimension contributed more weakly than the horizontal dimension did to the EMREO signal in two monkeys (U, J), but more strongly in a third (Y); the two dimensions exerted a roughly comparable effect in the fourth (F). Similarly, while some subjects have clear, significant oscillations in the constant component (Monkey F and Monkey U), others have a much noisier signal (Monkey Y) (Figure 2b bottom row). The constant component of the monkeys also differs from the humans in that the largest positive peaks occur later in the time course of the EMREO (around 75 ms-100 ms) compared to within the first 50 ms after saccade onset in humans.

A population level comparison across all subject ears is illustrated in Figure 3A-C. This figure shows the average of all human data for both the visually-guided saccade task (purple) and the free-viewing saccade task (green) as well as the monkey data for the free-viewing saccade task (black), aligned to onset of the saccade. The additional human subjects tested only on the visually-guided saccade task (n=17 subjects, n=34 ears) are also included (orange traces). As suggested in Figure 2, task has little effect on the EMREO signal within the same subjects (green vs. purple), and the results in this small sample accord well with the larger group tested only with the visual task (orange).

**Figure 3:**
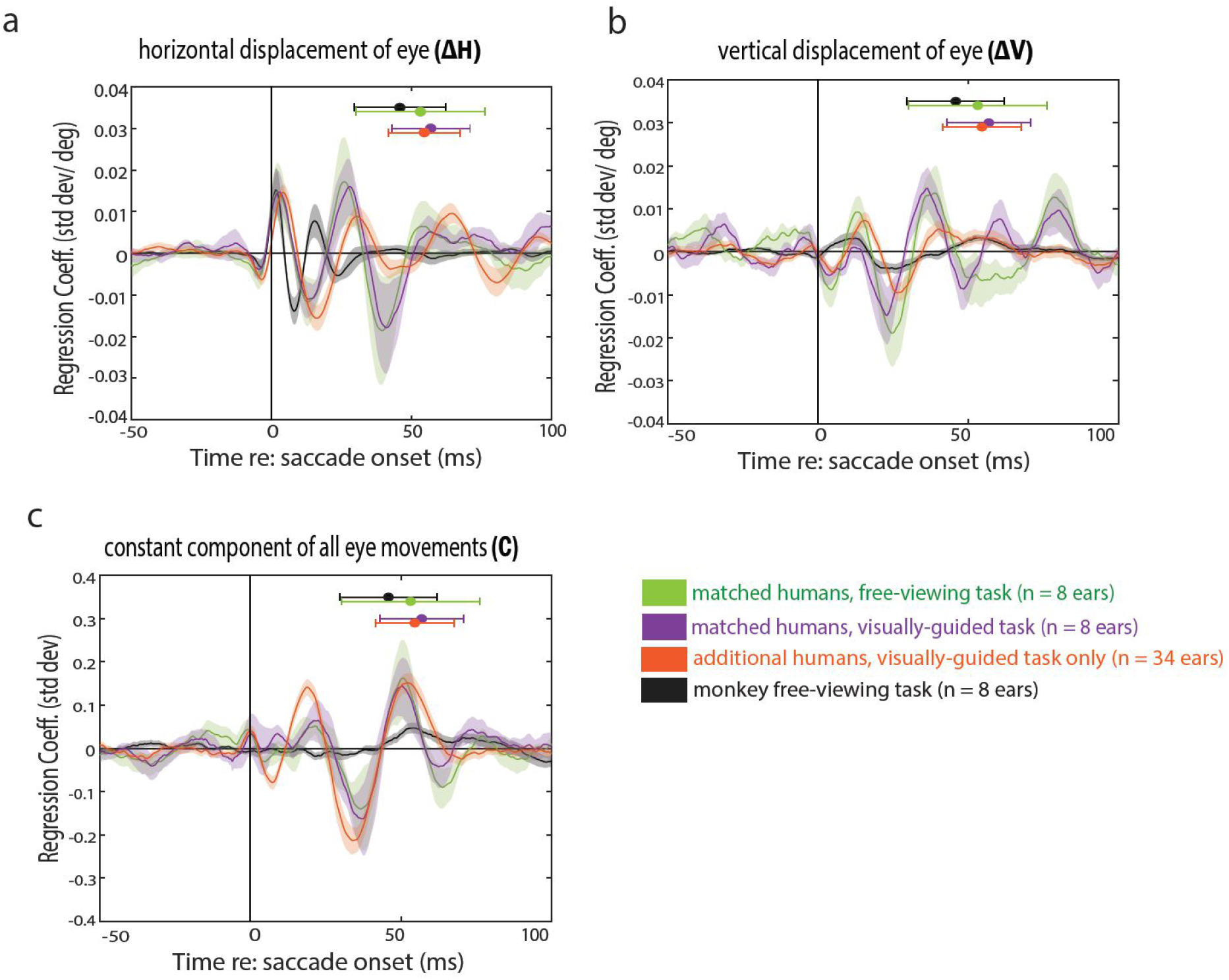
Average EMREO signal across populations of subjects. Average EMREO regression coefficients across both left and right ears are shown for 3 groups/sets of conditions for human subjects in comparison to the monkeys. Purple and green traces show the same set of 4 human participants as illustrated in figure 2, who performed the visually-guided saccade task (purple) and the free-viewing saccade paradigm (green). Orange traces show an additional larger group (N=17 subjects, 34 ears) who only performed the visually-guided saccade task. The monkey free-viewing saccades data are shown in black. The regression coefficient concerning horizontal displacement of the eye is plotted in (a), the coefficient for vertical displacement is shown in (b), and the constant component is plotted in (c). Regression coefficients were first computed within each subject ear. To facilitate combining across ears, the sign convention for the horizontal dimension was defined relative to the recorded ear (e.g. saccade amplitude was assigned a positive value for contralateral saccades and a negative value for ipsilateral saccades). The regression coefficients for each subject ear were then averaged together across all the relevant subjects (monkeys, human free viewing, corresponding human visually guided task, and the additional human subjects tested in visually guided tasks only). Shading indicates SEM across the subject ears included in each group. Mean saccade ending time +/- standard deviation for each species and task is plotted above the regression coefficients along the same time axis.

On average, the chief difference between the horizontal displacement regression coefficients in human vs. monkeys was in the waveform shape and time course. The oscillation initially occurs at a higher frequency for the monkey responses than the humans, and this rapid oscillatory portion then transitions to several much slower peaks/troughs.

The chief differences for the average vertical and constant components in humans vs. monkeys was in their magnitude (Figure 3b). For the constant component, the human data exhibits clear significant peaks and aligns in phase at the onset of the saccade for both tasks (purple: visually guided; green: free viewing). There is no comparably consistent signal in the monkey data. As seen in Figure 2, some individual monkeys (Monkey F and Monkey U) have significant positive and negative peaks between 50-100 ms, very different from what we see in the humans. However, the timing and waveform of these signals differed in these two animals, and the other two monkeys had no peaks in the constant component that were significantly different from zero. Therefore the average across the monkeys is much smaller than for the humans.

Similarly, even though all four individual monkeys exhibited vertical components, they were all different from each other in waveform, whereas in the humans this signal was not necessarily larger but its timing and waveform was more consistent across individuals. The net result is that when averaging across individual subjects, the vertical signal actually appears smaller in the monkeys than in the humans. It is unclear why the constant and vertical components show this inter-individual variability within monkeys or variability between the monkey and human groups.

To a degree, the faster time course of the monkey EMREO compared to humans may relate to the faster time course of monkey saccades combined with the known phase-resetting associated with saccade offset. As shown in Figure 3a-b, the saccades of the monkeys were shorter in duration than those of the humans (black vs. purple vs. green vs. orange bars at the top of the graphs). It is known that monkeys make faster saccades than humans (compare findings in: (Bahill et al., 1975; Jay and Sparks, 1990; Groh and Sparks, 1996; Guadron et al., 2022)), and this was true in our participants as well.

In sum, our findings indicate that the dependence of the EMREO on the horizontal displacement of the eye movement is quite conserved across humans and monkeys, differing mainly in the time course of the oscillation. Greater differences are observed in the vertical and constant components, and whether these signals appear comparable vs smaller in monkeys vs humans depends on whether the analysis is conducted at the level of individual participants or is averaged across the population.

## Discussion

Sounds and visual stimuli are detected in different “native” coordinate systems. The location of a sound is computed from cues involving loudness and timing differences across ears (interaural loudness differences or ILDs; interaural timing differences or ITDs), as well as variation in spectral frequency as a function of location with respect to the ears. These cues yield location information in a head-centered reference frame. The location of a visual stimulus is ascertained from the pattern of light hitting the retina, producing cues in an eye-centered reference frame. Information about the position and movements of the eyes with respect to the head is therefore necessary before the locations of visual and auditory stimuli can be matched up with one another. Understanding how the brain incorporates this information to reconcile these different coordinate systems is critical to knowing how the brain links information across the visual and auditory systems.

Research into the intersection between eye movements and hearing has involved psychophysical studies in humans (e.g. (Weerts and Thurlow, 1971; Lewald and Ehrenstein, 1996; Lewald, 1997, 1998; Boucher et al., 2001; Lewald and Ehrenstein, 2001; Lewald and Getzmann, 2006; Pavani et al., 2008; Klingenhoefer and Bremmer, 2009; Collins et al., 2010; Krüger et al., 2016; Willett et al., 2019)) and neurophysiological studies in monkeys (e.g. (Jay and Sparks, 1984, 1987b, a; Russo and Bruce, 1994; Hartline et al., 1995; Stricanne et al., 1996; Cohen and Andersen, 2000; Groh et al., 2001; Werner-Reiss et al., 2003; Fu et al., 2004; Zwiers et al., 2004; Mullette-Gillman et al., 2005; Porter et al., 2006; Mullette-Gillman et al., 2009; Maier and Groh, 2010; Bulkin and Groh, 2012b, a; Lee and Groh, 2012; Willett et al., 2019; Caruso et al., 2021); (See also: (Zella et al., 2001; Populin et al., 2004). The discovery of EMREOs in both humans and monkeys (Gruters et al., 2018; Murphy et al., 2020; Lovich et al., 2022; Abbasi et al., 2023; Bröhl and Kayser, 2023; King et al., 2023) afforded an opportunity to study an early aspect of this process in the same way in two different species. Given that how EMREOs actually contribute to auditory coordinate transformations has not yet been established, a comparison between human and monkey EMREOs provides an opportunity to identify clues as to which aspects of the EMREO signal are conserved across species.

We report here that in both humans and monkeys, EMREO waveforms depend on the magnitude and direction of the accompanying saccade, especially in the horizontal dimension. Performance of a visual task is not necessary; the signal is similar regardless of whether there is an explicit reason for making a saccade or if the participant is free-viewing. EMREOs occur without the presentation of visual or auditory stimuli and across an assortment of target locations, saccade magnitudes, and saccade tasks. The EMREO’s dependence on horizontal eye displacement exhibits a higher oscillation frequency in monkeys than humans, but a similar pattern is not seen for the vertical dimension, and the significance of this aspect of the signal is therefore unclear. A portion of this observation may simply reflect the fact that phase-resetting of the EMREO occurs at saccade offset, and monkey saccades are shorter in duration than human saccades. Put another way, if the period of the EMREO oscillation is roughly proportional to saccade duration, the systematic differences in saccade duration between monkeys and humans could account for this finding. However, it is unclear why a similar pattern is not clearly observed in the other aspects of the EMREO such as the vertical and constant components. Differences in teh resonance properties of the eardrum or associated structures could also contribute to these frequency/shape differences.

In contrast to the largely similar horizontal displacement component (within and across species), the waveforms and amplitudes of the vertical and constant components in the EMREO regression analysis differed across humans and monkeys, and, in the latter, across subjects. The constant term reflects the average EMREO waveform across all directions and amplitudes of movement. In humans, there appears to be a basic non-zero EMREO waveform upon which the dependence on eye movement parameters is superimposed. In monkeys, this basic EMREO waveform is much smaller and variable across individuals. The vertical signal is also sufficiently variable across monkeys that it largely disappears when averaging at the population level (Figure 3), leaving chiefly the dependence of the waveform on the eye movement parameters as the dominant signal. That the basic EMREO waveform is different between humans and monkeys, and shows different properties in the horizontal vs. vertical dimensions despite largely similar visual-auditory-oculomotor constraints and performance in the two species, suggests that the overall shape of the EMREO waveform may be of less significance than its dependence on eye movement parameters. As long as the signal is reproducible within an individual participant, it may contain the signatures needed for the brain to interpret its hypothesized impact on sound transduction in an eye movement–dependent fashion.

Our study provides an advance over our previous work (Gruters et al., 2018) in several ways. Our previous study presented monkey data in aggregated form only, and while the results appeared qualitatively similar to the aggregated human data, we did not perform an analysis of individual monkey data to establish that parallel on a more formal basis. Here, we used the same regression analysis on both human and monkey data, and we evaluated the pattern of results at both the individual and population levels. We also focused on both the dependence of the EMREO on the horizontal and vertical displacements of the eyes as well as the average EMREO waveform for comparison purposes.

Our study has several limitations. The free-viewing method did not permit fully exploring aspects of the EMREO signal relating to the initial position of the eyes. In addition, the free viewing paradigm may not have yielded fully comparable distributions of eye movements across individuals and species, and this could have led to different patterns of “variance capture” by the different regression terms across individuals and species. The lack of a visually-guided saccade task also limits the accuracy of our eye tracking calibration. Future work with a controlled task will be needed to clarify these points. A second related limitation of both the current work and our previous work is that the use of a microphone to measure the efferent system of the ear limits the analysis to oscillatory aspects of the signal; it is possible that there are steady state changes that are not reflected in the measurements made here. Such steady state changes can potentially be detected via laser doppler vibrometry or optical coherence tomography (Starovoyt et al., 2019) and may provide more information about sensitivity to initial/final eye position.

A third limitation is that we did not attempt to quantify the overall amplitude of the EMREO in humans vs. monkeys. This would have been challenging for several reasons. The microphone used here is specialized for the frequency range of conventional otoacoustic emissions (e.g. > 300 Hz), and thus its gain in the much lower frequency range of EMREOs is less certain (we have previously estimated that the amplitude of the human EMREO for an 18 degree eye movement to be about 57 dB SPL, based on modeling of the microphone’s transfer function in this frequency range (Gruters et al., 2018). Second, the relationship between the microphone’s output and the amplitude of the eardrum oscillation will depend on the size and acoustic properties of the ear canal space, which likely differ between humans and monkeys. Accordingly, the microphone measurements used in this study are expressed in units of standard deviation relative to a baseline period. This permits comparison across species on an even footing, although it leaves uncertain the precise conversion to sound pressure level.

Our overarching theory is that the mechanism(s) underlying EMREOs contribute to the process of localizing an incoming sound with respect to the eyes. How exactly this may happen remains uncertain. The cues to sound location involve interaural timing differences, interaural level differences, and the spectral content of sound. Eye-movement-related adjustments to any of these cues could contribute to the computation of sound location with respect to the eyes for subsequent integration with visual signals.

Understanding how this might occur requires considering the potential role(s) of the ear’s motor elements in generating EMREOs. These candidate motor elements consist of the two middle ear muscles (MEMs), the stapedius and tensor tympani, and the outer hair cells (OHCs). Generally, all three of these motor elements are thought to control the gain of the response of the ear to sound, with the MEMs thought of as playing a role in dampening the responses to loud sounds and the OHCs thought of as enhancing the responses to quiet sounds. Adjusting the gain of the response *differently* in the two ears in an eye movement dependent fashion would likely affect the inferred interaural level difference cues. For example, if deviation of the eyes to the left were to cause the gain of sound transduction to decrease in the left ear and/or increase in the right ear, the resulting alteration in the binaural level difference produced by incoming sounds would no longer be anchored solely to the head but could then be interpreted as indicating sound location with respect to the eyes. In addition, recent work by (Cho et al., 2023) has suggested that contractions of the middle ear muscles should not only influence the gain of responsiveness to incoming sound, but also the time delay of sound transmission, and thus potentially interaural timing difference cues in a similar fashion. Both timing and level difference cues relate solely to the horizontal dimension, which is the most consistent and powerful contributor to the EMREO signal in both humans and monkeys. How spectral cues are processed in the brain is less well known, but the ability of OHCs to alter processing in a frequency specific fashion could be relevant to how this cue might be modulated in an eye movement dependent way.

Candidate anatomical pathways by which oculomotor commands could reach any or all of these actuators exist. For example, the superior colliculus, a brain region involved in controlling eye movements, projects to the inferior colliculus (IC), an auditory structure (Coleman and Clerici, 1987; Sparks and Hartwich-Young, 1989). Auditory signals in both the SC and the IC are sensitive to changes in eye position (SC: (Jay and Sparks, 1984, 1987b, a; Hartline et al., 1995; Zella et al., 2001; Populin et al., 2004; Lee and Groh, 2012) ; IC: (Groh et al., 2001; Zwiers et al., 2004; Porter et al., 2006; Bulkin and Groh, 2012a, b; Willett et al., 2019), and the IC conveys descending projections to both the superior olivary complex (Faye-Lund, 1986), the source of descending input to the outer hair cells (Guinan, 2006; Ciuman, 2010) and the cochlear nucleus (Milinkeviciute et al., 2017; Balmer and Trussell, 2022) which in turn projects to the facial and trigeminal nerves that innervate the stapedius and tensor tympani respectively (Mukerji et al., 2010). Preliminary findings from our group are consistent with a role for all three of these types of motor actuators in the generation of EMREOs (King et al., 2023). Future studies regarding these motor components and how they control mechanical processes in the ear will benefit from an animal model with a significant and consistent EMREO and with comparable ear anatomy to humans. With an animal model similar enough to humans, interventional studies can be conducted to isolate the source of motor commands and their impact on the motor actuators that create and/or modulate the EMREO signal.

Overall, it is important to understand the similarities and differences of the EMREO signal across species when considering future work to determine which anatomical components are necessary and functional in the EMREO signal. With information about the key features that are conserved in the signal, we can answer questions about the physical mechanisms in the ear that generate the EMREO and lead to sound localization and, eventually, perception.

## References

Abbasi H, King CD, Lovich S, Röder B, Groh JM, Bruns P (2023) Audiovisual temporal recalibration modulates eye movement-related eardrum oscillations. International Multisensory Research Forum.

Bahill AT, Clark MR, Stark L (1975) The main sequence, a tool for studying human eye movements. Mathematical Biosciences 24:191–204.

Balmer TS, Trussell LO (2022) Descending Axonal Projections from the Inferior Colliculus Target Nearly All Excitatory and Inhibitory Cell Types of the Dorsal Cochlear Nucleus. 42:3381–3393.

Boucher L, Groh JM, Hughes HC (2001) Afferent delays and the mislocalization of perisaccadic stimuli. Vision Research 41:2631–2644.

Bröhl F, Kayser C (2023) Detection of spatially-localized sounds is robust to saccades and concurrent eye movement-related eardrum oscillations (EMREOs). BioRxiv:2023.2004.2017.537161.

Bulkin DA, Groh JM (2012a) Distribution of visual and saccade related information in the monkey inferior colliculus. Front Neural Circuits 6:61.

Bulkin DA, Groh JM (2012b) Distribution of eye position information in the monkey inferior colliculus. J Neurophysiol 107:785–795.

Caruso VC, Pages DS, Sommer MA, Groh JM (2019) Compensating for a shifting world: A quantitative comparison of the reference frame of visual and auditory signals across three multimodal brain areas. bioRxiv:669333.

Caruso VC, Pages DS, Sommer MA, Groh JM (2021) Compensating for a shifting world: evolving reference frames of visual and auditory signals across three multimodal brain areas. J Neurophysiol 126:82–94.

Cho NH, Ravicz ME, Puria S (2023) Human middle-ear muscle pulls change tympanic-membrane shape and low-frequency middle-ear transmission magnitudes and delays. Hear Res 430:108721.

Ciuman RR (2010) The efferent system or olivocochlear function bundle - fine regulator and protector of hearing perception. International journal of biomedical science : IJBS 6:276–288.

Cohen YE, Andersen RA (2000) Reaches to sounds encoded in an eye-centered reference frame. Neuron 27:647–652.

Coleman JR, Clerici WJ (1987) Sources of projections to subdivisions of the inferior colliculus in the rat. J Comp Neurol 262:215–226.

Collins T, Heed T, Roder B (2010) Eye-movement-driven changes in the perception of auditory space. Attention, perception & psychophysics 72:736–746.

Faye-Lund H (1986) Projection from the inferior colliculus to the superior olivary complex in the albino rat. Anatomy and embryology 175:35–52.

Fu KM, Shah AS, O’Connell MN, McGinnis T, Eckholdt H, Lakatos P, Smiley J, Schroeder CE (2004) Timing and laminar profile of eye-position effects on auditory responses in primate auditory cortex. J Neurophysiol 92:3522–3531.

Groh JM (2014) Your sunglasses are in the milky way. In: Making space: how the brain knows where things are. Cambridge, MA: Harvard University Press.

Groh JM, Sparks DL (1992) Two models for transforming auditory signals from head-centered to eye-centered coordinates. Biological Cybernetics 67:291–302.

Groh JM, Sparks DL (1996) Saccades to somatosensory targets. I. behavioral characteristics. J Neurophysiol 75:412–427.

Groh JM, Trause AS, Underhill AM, Clark KR, Inati S (2001) Eye position influences auditory responses in primate inferior colliculus. Neuron 29:509–518.

Gruters KG, Murphy DLK, Jenson CD, Smith DW, Shera CA, Groh JM (2018) The eardrums move when the eyes move: A multisensory effect on the mechanics of hearing. Proc Natl Acad Sci U S A 115 E1309–E1318.

Guadron L, van Opstal AJ, Goossens J (2022) Speed-accuracy tradeoffs influence the main sequence of saccadic eye movements. Scientific reports 12:5262.

Guinan JJ, Jr. (2006) Olivocochlear efferents: anatomy, physiology, function, and the measurement of efferent effects in humans. Ear Hear 27:589–607.

Hartline PH, Vimal RL, King AJ, Kurylo DD, Northmore DP (1995) Effects of eye position on auditory localization and neural representation of space in superior colliculus of cats. Exp Brain Res 104:402–408.

Jay MF, Sparks DL (1984) Auditory receptive fields in primate superior colliculus shift with changes in eye position. Nature 309:345–347.

Jay MF, Sparks DL (1987a) Sensorimotor integration in the primate superior colliculus. II. Coordinates of auditory signals. J Neurophysiol 57:35–55.

Jay MF, Sparks DL (1987b) Sensorimotor integration in the primate superior colliculus. I. Motor convergence. J Neurophysiol 57:22–34.

Jay MF, Sparks D (1990) Localization of auditory and visual targets for the initiation of saccadic eye movements. In: Comparative Perception Vol I Basic Mechanisms (Berkley MA, Stebbins WC, eds), p 527. New York: John Wiley & Sons.

Kemp DT (1978) Stimulated acoustic emissions from within the human auditory system. The Journal of the Acoustical Society of America 64:1386–1391.

King CD, Lovich SN, Murphy DL, Landrum R, Kaylie D, Shera CA, Groh JM (2023) Individual similarities and differences in eye-movement-related eardrum oscillations (EMREOs). BioRxiv:2023.2003.2009.531896.

King CD, Lovich SNS, Murphy DLK, Landrum R, Kaylie DM, Shera CA, Groh JM (submitted) Individual similarities and differences in eye-movement-related eardrum oscillations (EMREOs).

Klingenhoefer S, Bremmer F (2009) Perisaccadic localization of auditory stimuli. Exp Brain Res 198:411–423.

Krüger HM, Collins T, Englitz B, Cavanagh P (2016) Saccades create similar mislocalizations in visual and auditory space. 115:2237–2245.

Lee J, Groh JM (2012) Auditory signals evolve from hybrid- to eye-centered coordinates in the primate superior colliculus. J Neurophysiol 108:227–242.

Lewald J (1997) Eye-position effects in directional hearing. Behavioural Brain Research 87:35–48.

Lewald J (1998) The effect of gaze eccentricity on perceived sound direction and its relation to visual localization. Hear Res 115:206–216.

Lewald J, Ehrenstein WH (1996) The effect of eye position on auditory lateralization. Experimental Brain Research 108:473–485.

Lewald J, Ehrenstein WH (2001) Effect of gaze direction on sound localization in rear space. Neuroscience Research 39:253–257.

Lewald J, Getzmann S (2006) Horizontal and vertical effects of eye-position on sound localization. Hear Res 213:99–106.

Lovich SN, King CD, Murphy DL, Landrum R, Shera CA, Groh JM (2022) Parametric information about eye movements is sent to the ears.2022.2011.2027.518089.

Maier JX, Groh JM (2010) Comparison of gain-like properties of eye position signals in inferior colliculus versus auditory cortex of primates. Frontiers in Integrative Neuroscience 4:121–132.

Milinkeviciute G, Muniak MA, Ryugo DK (2017) Descending projections from the inferior colliculus to the dorsal cochlear nucleus are excitatory. 525:773–793.

Mohl JT, Pearson JM, Groh JM (2019) Monkeys and humans implement causal inference to simultaneously localize auditory and visual Stimuli. BioRxiv; Submitted, PLOS Computational Biology.

Mukerji S, Windsor AM, Lee DJ (2010) Auditory brainstem circuits that mediate the middle ear muscle reflex. Trends Amplif 14:170–191.

Mullette-Gillman OA, Cohen YE, Groh JM (2005) Eye-centered, head-centered, and complex coding of visual and auditory targets in the intraparietal sulcus. J Neurophysiol 94:2331–2352.

Mullette-Gillman OA, Cohen YE, Groh JM (2009) Motor-related signals in the intraparietal cortex encode locations in a hybrid, rather than eye-centered, reference frame. Cerebral Cortex 19:1761–1775.

Murphy DL, King CD, Schlebusch SN, Landrum R, Shera CA, Groh JM (2020) Evidence for a system in the auditory periphery that may contribute to linking sounds and images in space. bioRxiv:2020.2007.2019.210864.

Pavani F, Husain M, Driver J (2008) Eye-movements intervening between two successive sounds disrupt comparisons of auditory location. Exp Brain Res 189:435–449.

Populin LC, Tollin DJ, Yin TC (2004) Effect of eye position on saccades and neuronal responses to acoustic stimuli in the superior colliculus of the behaving cat. J Neurophysiol 92:2151–2167.

Porter KK, Metzger RR, Groh JM (2006) Representation of eye position in primate inferior colliculus. J Neurophysiol 95:1826–1842.

Russo GS, Bruce CJ (1994) Frontal eye field activity preceding aurally guided saccades. J Neurophysiol 71(3):1250–1253.

Schairer KS, Feeney MP, Sanford CA (2013) Acoustic Reflex Measurement. 34:43s–47s.

Shera CA (2004) Mechanisms of mammalian otoacoustic emission and their implications for the clinical utility of otoacoustic emissions. Ear Hear 25:86–97.

Sparks DL, Hartwich-Young R (1989) The deep layers of the superior colliculus. In: The neurobiology of saccadic eye movements (Wurtz RH, Goldberg ME, eds), pp 213–255. New York: Elsevier.

Starovoyt A, Putzeys T, Wouters J, Verhaert N (2019) High-resolution Imaging of the Human Cochlea through the Round Window by means of Optical Coherence Tomography. Scientific reports 9:14271.

Stricanne B, Andersen RA, Mazzoni P (1996) Eye-centered, head-centered, and intermediate coding of remembered sound locations in area LIP. J Neurophysiol 76:2071–2076.

Weerts TC, Thurlow WR (1971) The effects of eye position and expectation on sound localization. Perception & Psychophysics 9:35–39.

Werner-Reiss U, Kelly KA, Trause AS, Underhill AM, Groh JM (2003) Eye position affects activity in primary auditory cortex of primates. Current Biology 13:554–562.

Willett SM, Groh JM, Maddox RK (2019) Hearing in a “moving” visual world: Coordinate transformations along the auditory pathway. In: Springer Handbook of Auditory Research: Multisensory Processes The Auditory Perspective (Lee AKC, Wallace MT, Coffin AB, Popper AN, Fay RR, eds): Springer.

Zella JC, Brugge JF, Schnupp JW (2001) Passive eye displacement alters auditory spatial receptive fields of cat superior colliculus neurons. Nat Neurosci 4:1167–1169.

Zwiers MP, Versnel H, Van Opstal AJ (2004) Involvement of monkey inferior colliculus in spatial hearing. J Neurosci 24:4145–4156.

